# Click chemistry selectively activates an auristatin protodrug with either intratumoral or systemic tumor-targeting agents

**DOI:** 10.1101/2023.03.28.534625

**Authors:** Jesse M. McFarland, Maša Alečković, George Coricor, Sangeetha Srinivasan, Matthew Tso, John Lee, Tri-Hung Nguyen, José M. Mejía Oneto

**Affiliations:** Shasqi Inc., 665 3rd St, Suite 501, San Francisco, CA 94107, USA

**Keywords:** Click Activated Protodrugs Against Cancer (CAPAC), click chemistry, local drug activation, oncology, prodrug

## Abstract

The Click Activated Protodrugs Against Cancer (CAPAC™) platform enables activation of powerful cancer drugs at tumor sites, maximizing efficacy and minimizing systemic toxicity. CAPAC utilizes a potent click chemistry reaction between tetrazine and *trans*-cyclooctene, called tetrazine ligation. The reaction between the activator, linked to a tumor targeting agent, and the protodrug leads to targeted activation of the drug.

In this study, activation is accomplished either by intratumoral injection of a tetrazine-modified hyaluronic acid biopolymer (SQL70) or by systemic infusion of a tetrazine-modified HER2-targeting antigen-binding fragment (SQT01). The drug used is monomethyl auristatin E (MMAE), a cytotoxic agent hindered in its clinical use by severe toxicity.

MMAE modification with a *trans*-cyclooctene moiety to form the protodrug SQP22 reduced its cytotoxicity *in vitro* and *in vivo*. Treatment of SQP22 paired with SQL70 biopolymer demonstrated anti-tumor effects in Karpas 299 and RENCA murine tumor models, establishing the requirement of click chemistry for protodrug activation. Furthermore, SQP22 paired with SQT01 induced anti-tumor effects in the HER2-positive NCI-N87 murine tumor model, showing that activation could be accomplished by systemic dosing of a tumor-targeting antibody conjugate. Observed toxicities were limited to modest, transient myelosuppression and moderate body weight loss in these tumor models.

This study further delineates the capabilities of the CAPAC platform by demonstrating anti-tumor activity of SQP22 with two differentiated targeting approaches and underscores the power of click chemistry to precisely control the activation of cancer drugs at tumor sites.

**TOC graphic:** 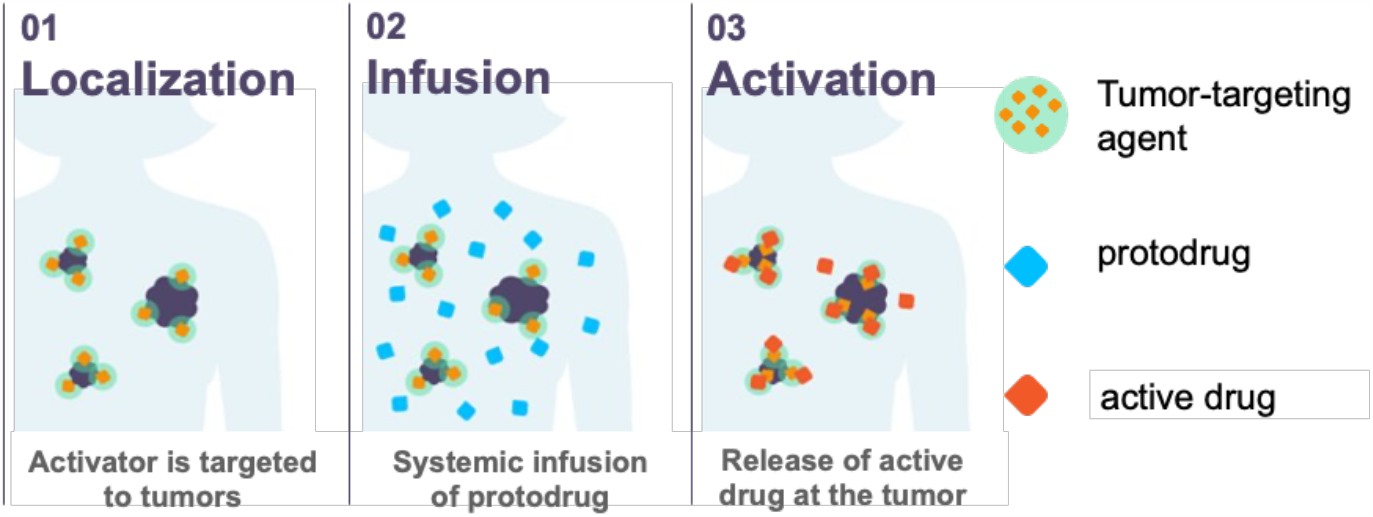

**Synopsis:** Click chemistry efficiently activates protodrugs at tumors. A novel protodrug delivering monomethyl auristatin E, **SQP22**, and preclinical experiments with a biopolymer and antibody fragment as targeting agents are reported.

## Introduction

Click chemistry encompasses chemical reactions that are fast, efficient, and selective in complex environments.^1^ The 2022 Nobel Prize in Chemistry was awarded to Sharpless, Meldal, and Bertozzi in recognition of the transformative effect click and bioorthogonal chemistry have had in research and drug development.^2^ The Click Activated Protodrugs Against Cancer (CAPAC™) platform developed by Shasqi (San Francisco, CA, USA) is an approach that uses chemistry to engineer biology and precisely control the activation of drugs at tumor sites.^3^ Specifically, the CAPAC platform uses the tetrazine ligation reaction between tetrazine and *trans*-cyclooctene,^4^ an exceptionally fast and specific click chemistry reaction compatible with biological environments.^5-11^

Cancer is a major challenge worldwide, with estimations of 20 million new cancer cases and 10 million cancer-related deaths in 2020.^12^ Advances in targeted therapies, including immunotherapies, biomarker-targeting therapies, and antibody drug conjugates (ADCs), have revolutionized the treatment of certain cancers. Unfortunately, only subsets of patients expressing sufficient levels of specific biomarkers benefit from targeted approaches.^13-16^ This leaves systemically administered cytotoxic chemotherapeutic agents as a standard-of-care for a wide variety of cancers despite their severe, often dose-limiting adverse effects and narrow therapeutic windows. The objective of the CAPAC platform is to address the clear need for an effective alternative to activate cancer therapies at tumors in the body.

The first investigational therapy based on click chemistry in humans, SQ3370, consists of two components: a tetrazine-modified biopolymer targeting agent (SQL70 biopolymer), which is injected intratumorally, and a *trans*-cyclooctene-modified protodrug of doxorubicin (SQP33 protodrug), which is infused intravenously (IV).^17^ The efficient *in vivo* reaction between the tetrazine moiety of SQL70 and the *trans*-cyclooctene moiety of SQP33 activates the protodrug, leading to release of active doxorubicin at the tumor site. Results from the dose escalation in a phase 1/2a, first-in-human, single-arm, open-label trial in advanced solid tumors (<u>NCT04106492</u>) showed that SQ3370 is safe, well tolerated and induces a cytotoxic T-cell supportive tumor microenvironment,^18,19^ consistent with data from previously published animal studies.^17^

Monomethyl auristatin E (MMAE) is a synthetic analog of the natural product dolastatin 10 and a potent anti-mitotic agent that inhibits tubulin polymerization.^20,21^ MMAE has 100–1,000 times more potent anti-tumor activity than doxorubicin; however, its use has been hindered by severe toxicities, including myelosuppression and neuropathy.^22,23^ Researchers have investigated several approaches to precisely target MMAE to tumors as a way to reduce its toxicity, including developing peptide conjugates^22,24^ and ADCs.^16,25,26^ The vedotin payload developed by Seagen Inc (Bothell, WA, USA) links MMAE to antibodies via a protease cleavable linker. Four vedotin ADCs have been approved by the US Food and Drug Administration (FDA) for cancer treatment: brentuximab vedotin,^27^ enfortumab vedotin-ejfv,^28^ polatuzumab vedotin-piiq,^29^ and tisotumab vedotin-tftv,^30^ validating the benefits of MMAE payloads in oncology. To activate and release the MMAE payload, internalization of the vedotin conjugates into tumor cells followed by processing by endogenous proteases is required. This limits the scope of antigens amenable to targeting by vedotin ADCs. Other conditional activation strategies rely on physiological factors such as tumor-specific biomarkers, enzymatic activity, pH, or reactive oxygen levels to activate the payload based on differences between tumors and healthy tissues.^16,25,26,31-35^ The common characteristic of these approaches is that they rely on activation by biological parameters and thus are limited by inherent variations within a single tumor and across patients.

The CAPAC platform, based on the tetrazine ligation reaction, was used to develop SQP22 (Figure 1A), a protodrug of MMAE, which is activated by tumor-targeting agents to release the MMAE payload specifically at tumors. Initially, we characterized SQP22 *in vitro* and *in vivo* paired with the SQL70 biopolymer^17^ (Figure 1B). SQL70 is injected intratumorally and activates multiple systemically administered SQP22 protodrug doses to release MMAE at the tumor (Figure 1C). To explore the activation of SQP22 *in vivo* without tumor injections, we developed a human epidermal growth factor receptor 2 (HER2) antigen-binding fragment (Fab) conjugated to tetrazine and designated it SQT01. By targeting a tumor-associated antigen, SQT01 can localize the tetrazine activators at the tumors that express the specific antigen. Desirable properties of the targeting antibody conjugate include: 1) tight binding to the tumor site with a slow off-rate, and 2) rapid clearance from circulation to minimize systemic activation of the infused protodrug.

**Figure 1.**
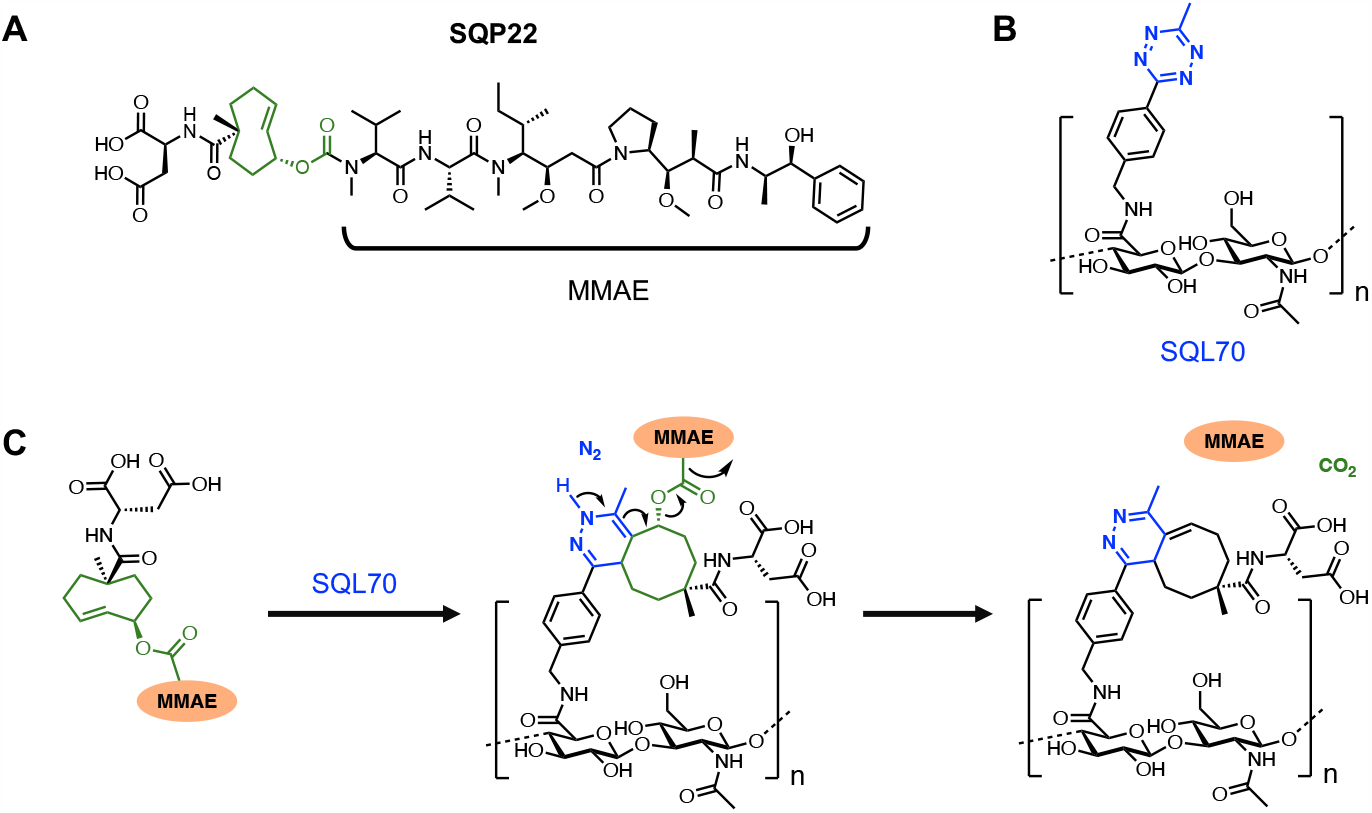
Activation of SQP22 protodrug by SQL70 biopolymer. **A**, Chemical structure of SQP22, a TCO-modified protodrug of MMAE. **B**, Chemical structure of Tz-modified biopolymer, SQL70. **C**, Schematic of the click chemistry reaction by which SQP22 is activated by Tz on SQL70 biopolymer and active MMAE is released. MMAE, monomethyl auristatin E; TCO, *trans*-cyclooctene; Tz, tetrazine.

In this article, we report the development of SQP22, an MMAE-based protodrug, and a companion tumor-targeting agent, the antibody-directed SQT01 conjugate. SQP22, when paired with either the intratumorally-injected SQL70 biopolymer or SQT01 elicited sustained anti-tumor responses in the Karpas 299, RENCA, and NCI-N87 murine tumor models. Materials and methods for this article are provided in Electronic Supporting Information.

## Results and discussion

### SQP22 cytotoxicity and stability in plasma and tissue homogenates

Modification of MMAE to form SQP22 protodrug reduced its cytotoxicity greater than 50-fold across several cell lines (Table 1). Activation of SQP22 by tetrazine activators led to the efficient release of MMAE and restored the activity of the drug with IC_50_ values in the 2-5 nM range. SQP22 was highly stable in plasma, with approximately 94% and 100% of the protodrug remaining after a 4-hour incubation period at 37 °C in human and mouse plasma, respectively (Table S1). Although up to 50% loss of SQP22 was observed when incubated over 24 hours in tissue homogenates at 37 °C, no MMAE was released, supporting its potential safety *in vivo* (Figure S1).

**Table 1.** *In vitro* cytotoxicity of SQP22

| **Cell Line** | **Cytotoxicity IC<sub>50</sub> (nM)** | **Fold Attenuation** |
| --- | --- | --- |
| MC38 | 5.5 | 704 |
| EMT6 | 2.8 | 133 |
| 4T1 | 3.8 | > 50 |
| B16-F10 | 2.7 | > 67 |
| RENCA | 1.8 | > 100 |

Cytotoxicity of SQP22 prodrug after activation with tetrazine is shown as IC<sub>50</sub> across several murine cancer cell lines. Fold attenuation was calculated based on the IC<sub>50</sub> values in the presence of tetrazine (activated drug) relative to the absence of tetrazine (attenuated prodrug).

### In vivo anti-tumor efficacy of SQP22 in various murine tumor models

#### SQP22 leads to complete tumor regression when combined with SQL70 in the Karpas 299 xenograft model

The attenuated cytotoxicity of SQP22 alone and its anti-tumor activity in the presence of SQL70 biopolymer were evaluated in the Karpas 299 xenograft model (Figure 2). While active MMAE administered as a single dose at 0.5 mg/kg (1x) led to a two-fold reduction in tumor burden compared to the vehicle control at day 15 (*P* < 0.0001), SQP22 in the absence of SQL70 biopolymer had no effect on tumor growth relative to the vehicle control (*P* = 0.62), even when administered as 5-daily doses of 10x molar equivalents of MMAE (50x cumulative dose) (Figure 2A, B). Furthermore, 1x MMAE resulted in about 6% body weight loss during dosing with body weights remaining significantly lower than in the vehicle control group at day 15 (*P* = 0.0011) (Figure 2C). On the other hand, SQP22 administered alone at 50x molar equivalents of MMAE led to minimal, transient body weight loss after dosing (< 3% on average) with no significant difference relative to the vehicle control at day 15 (*P* = 0.64), demonstrating efficient potency attenuation and safety of the protodrug.

**Figure 2.**
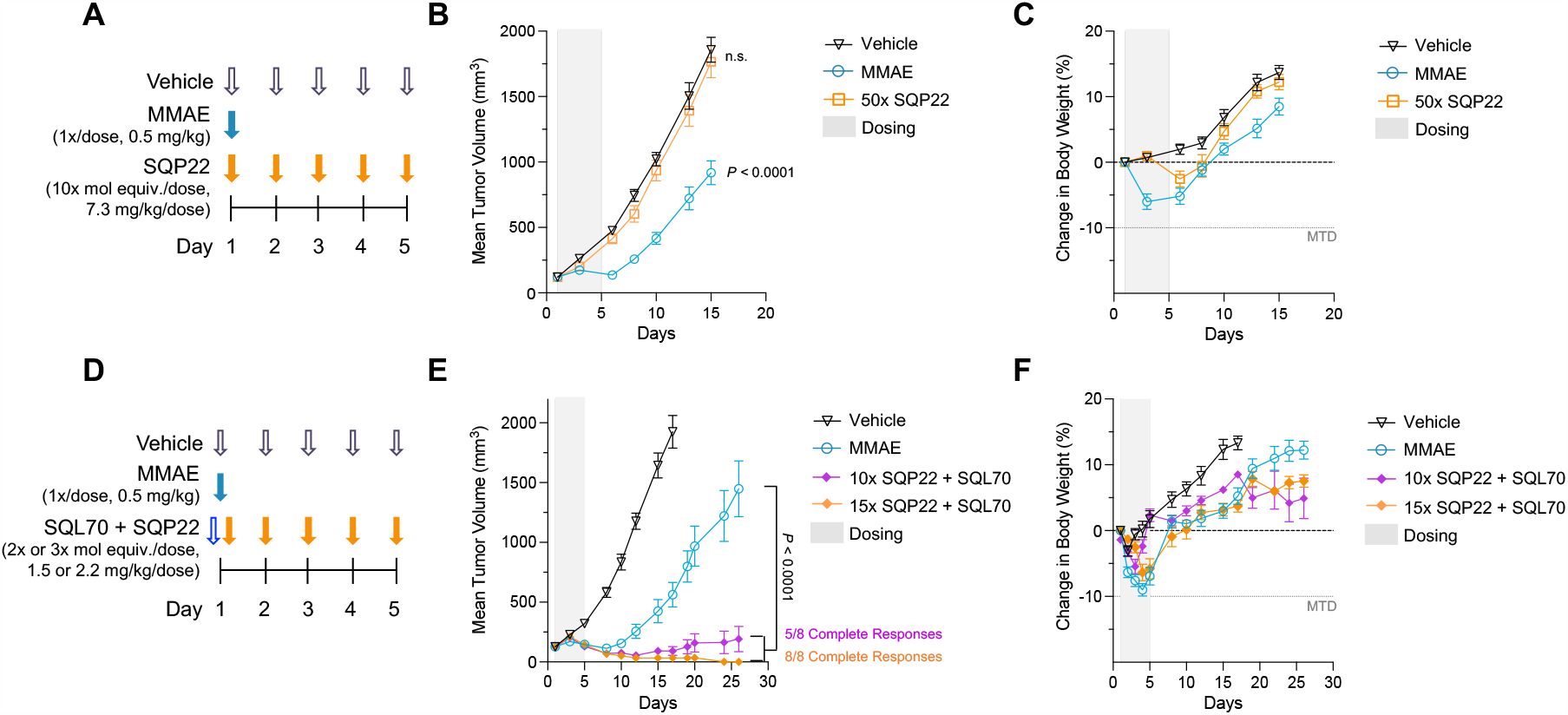
SQP22 leads to complete regression of Karpas 299 xenograft tumors in the presence of SQL70. **A**, Schedule of dosing of agents in absence of SQL70 biopolymer. **B-C**, Tumor volumes of Karpas 299 xenografts in C.B-17 SCID mice (**B**) and body weight change (**C**) after dosing with vehicle, MMAE, and SQP22 protodrug. Only MMAE led to significant body weight loss on day 15 (P < 0.0001). **D**, Schedule of dosing of agents in presence of SQL70. SQP22 was dosed 1 hour after SQL70 injection. **E-F**, Tumor volumes of Karpas 299 xenografts in C.B-17 SCID mice (**E**) and body weight change (**F**) after dosing with vehicle, MMAE, and SQP22 protodrug with the SQL70 biopolymer. Shown are means ± SEM (*n* = 8 mice/group). *P*-values were determined by two-way ANOVA with Bonferroni correction for multiple comparisons. Complete response was defined as no palpable tumors measured in 3 consecutive days. ANOVA, analysis of variance; MMAE, monomethyl auristatin E; mol equiv., molar equivalent; MTD, maximum tolerated dose; SEM, standard error of mean.

When administered in the presence of SQL70 biopolymer (Figure 2D), SQP22 at cumulative doses of 10x and 15x led to improved anti-tumor efficacy compared with 1x MMAE treatment starting on day 10, with eventual complete tumor regression in 5 of 8 and 8 of 8 animals, respectively (Figure 2E, Figure S2), suggesting dose-dependent effects of SQP22. MMAE alone did not lead to complete responses in tumor-bearing animals. This demonstrates significant enhancement in efficacy by SQP22 at either dose with SQL70 compared with MMAE treatment (day 26, *P* < 0.0001) (Figure 2E).

Body weight loss in the MMAE, 10x SQP22, and 15x SQP22 treatment groups was comparable and transient after dosing initiation, with the 10x SQP22 group recovering faster than the other groups (Figure 2F). Starting on day 20, the body weight change diverged between the 10x and 15x SQP22 treatment groups. Notably, body weight in the treated animals did not drop more than 10%, suggesting manageable treatment-induced toxicity.

#### SQP22 with SQL70 inhibits RENCA tumor progression with transient effects on complete blood count

The effect of SQP22 in combination with the targeting biopolymer, SQL70, was next evaluated in the RENCA syngeneic tumor model (Figure 3A). MMAE treatment at 1 mg/kg (1x) resulted in reduction of tumor growth with a four-fold difference in tumor volumes at day 16 (Figure 3B, day 16, *P* < 0.0001), supporting the use of the RENCA model as a MMAE-sensitive syngeneic tumor model. Mice administered only SQL70 biopolymer showed no difference in tumor volume (Figure 3B) or body weight change compared with mice administered vehicle control (Figure S3A). However, administration of SQL70 followed by SQP22 as 3 daily doses of 2x or 3x molar equivalents of MMAE (6x or 9x cumulative dose, respectively) led to significant reduction in tumor progression compared with administration of vehicle control (day 16, *P* < 0.0001 for both) or MMAE (day 30, *P* = 0.0075 and *P* < 0.0001, respectively), supporting the strong anti-tumor effects of SQP22 treatment even at a reduced dosing schedule. In the presence of SQL70, SQP22 showed greater anti-tumor efficacy at 9x compared with the 6x cumulative MMAE molar equivalents dose (*P* = 0.0008), confirming a dose response over the study duration.

**Figure 3.**
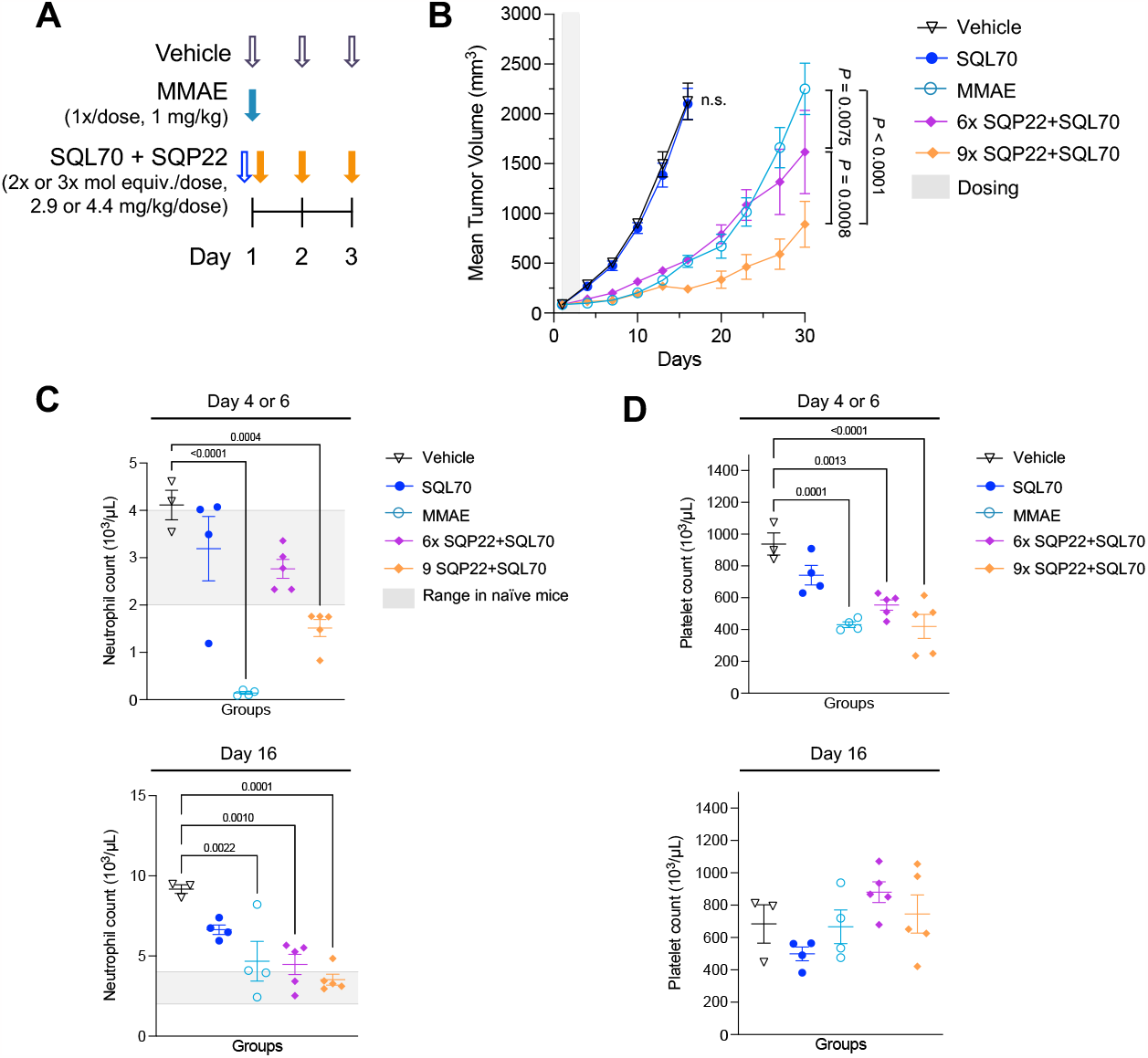
SQP22 with SQL70 reduces growth of RENCA syngeneic tumors with reversible effects on complete blood counts. **A**, Schedule of dosing of agents. SQP22 was dosed 1 hour after SQL70 injection. **B**, Tumor volumes of RENCA tumors in BALB/c mice following treatment with vehicle (*n* = 3 mice), MMAE (*n* = 4 mice), SQL70 alone (n = 4 mice), and SQL70 with SQP22 dosed at 2x (*n* = 5 mice) and 3x (*n* = 5 mice) mol equiv. of MMAE/dose. **C-D**, On the indicated days, blood samples were collected and analyzed for neutrophil (**C**) and platelet (**D**) counts. Range in neutrophil cell counts from naïve BALB/c mice is indicated by the grey box ^36^. Shown are means ± SEM. *P*-values were determined by two-way ANOVA with *post-hoc* Bonferroni correction for Day 30 (**B**) and one-way ANOVA with *post-hoc* Bonferroni correction (**C, D**). ANOVA, analysis of variance; MMAE, monomethyl auristatin E; mol equiv., molar equivalent; SEM, standard error of mean.

Of note, untreated RENCA-bearing mice lose body weight as tumors progress, hence we evaluated the acute effects of treatments by analyzing the maximum body weight change in the first week after dosing initiation. No difference in acute body weight loss was observed between mice that were administered SQL70 or vehicle control (Figure S3B). Unlike MMAE treatment, which resulted in ≥ 15% body weight loss within the first 7 days after dosing, neither dose of SQP22 in the presence of SQL70 resulted in a significant reduction in body weight compared with vehicle control (Figure S3A, B). In fact, acute body weight loss in mice treated with SQP22 and SQL70 was < 10%, consistent with results observed in the Karpas 299 model.

As MMAE has been shown to exhibit myelosuppressive effect, we next evaluated acute and longer-term changes in the complete blood count of the RENCA-bearing mice. Blood samples drawn 3 days after the final dose (day 4 for the MMAE group, day 6 for all other treatment groups) and on day 16 were analyzed for complete blood counts (Figure 3C, D). On day 4, acute neutropenia was observed in MMAE-treated mice relative to vehicle (*P <* 0.0001), while moderate (∼50%) neutrophil reduction was observed on day 6 in mice treated with SQP22 at the cumulative dose of 9x molar equivalents of MMAE in the presence of SQL70 relative to vehicle control (*P =* 0.0004). Neutrophils for both groups were below the normal range reported for wild-type BALB/c mice (i.e., 20–30% of white blood cell count).^36^ On the other hand, mice receiving the cumulative dose of 6x molar equivalents of MMAE displayed neutrophil counts within the normal range with no significant difference compared with the vehicle control group (*P* = 0.0653). Reductions in platelet counts to similar levels were measured in mice dosed with MMAE alone (*P* = 0.0001) and SQL70 with SQP22 at 6x (*P* = 0.0013) or 9x (*P* < 0.0001) cumulative doses (Figure 3D). Both acute effects observed, neutropenia and thrombocytopenia, were reversible by day 16, indicating transient MMAE-associated myelosuppression (Figure 3C, D).

#### SQP22 in combination with SQT01 inhibits tumor progression in the HER2-positive NCI-N87 xenograft model

SQT01 was designed as a Fab-tetrazine conjugate to balance the need to specifically bind the HER2 antigen with high affinity without the extended circulation associated with a full IgG. Preparation and characterization of SQT01 are presented in Electronic Supporting Information (Figure S4). Briefly, the purified Fab of trastuzumab was conjugated on lysine residues with tetrazine-PEG9-NHS and the resulting tetrazine-to-antibody ratio was determined to be 2.2. The conjugate was highly monomeric and bound antigen-positive cells similarly to the unconjugated Fab. A non-binding isotype control Fab-tetrazine conjugate was prepared and characterized by similar methods.

The effect of SQP22 in combination with SQT01 was evaluated in the NCI-N87 xenograft model (Figure 4A). While tumor-bearing mice treated with SQP22 alone displayed no effect on tumor growth compared with those treated with vehicle (*P* > 0.99), infusion of SQT01 4 hours prior to SQP22 treatment resulted in significant, five-fold reduction in tumor size compared with the vehicle controls on day 27 (Figure 4B, *P* < 0.0001). Moreover, SQP22 with SQT01 showed significant inhibition of tumor progression compared with SQP22 alone or SQP22 with the isotype control Fab-tetrazine conjugate (*P* < 0.0001). The minimal activity observed in the non-binding control group may be due to a small amount of the conjugate remaining in circulation at the time of SQP22 infusion. Disitamab vedotin is a HER2-targeted vedotin ADC being studied in multiple late-stage clinical trials targeting HER2-positive solid tumors.^37-39^ It was included as a positive control to benchmark the treatment against an ADC carrying an MMAE payload and showed only a two-fold reduction in tumor burden compared with vehicle at study end point (*P* < 0.0001). In fact, SQP22 with SQT01 showed superior efficacy compared with disitamab vedotin by day 27 (*P* = 0.0009).

**Figure 4.**
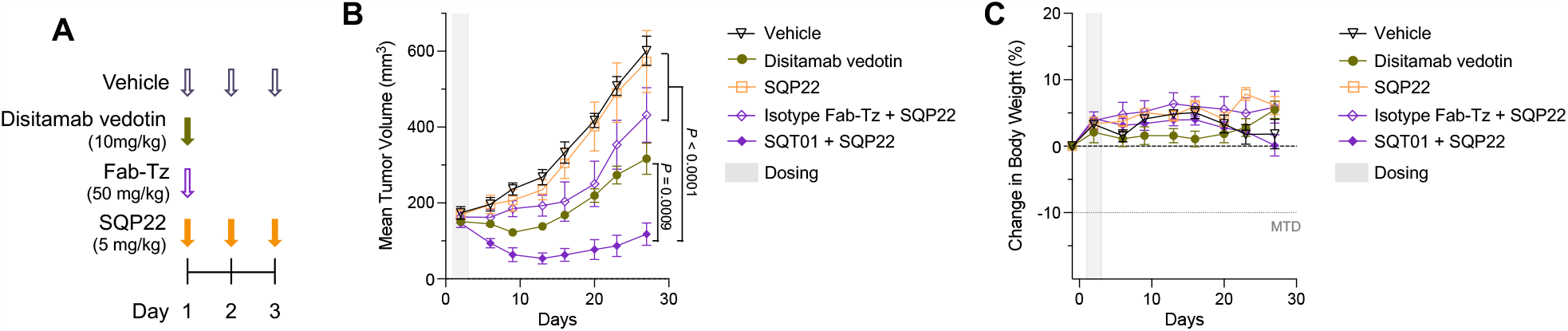
SQP22 leads to complete regression of NCI-N87 xenograft tumors in the presence of SQT01. **A**, Schedule of dosing of agents. Two types of Fab-Tz agents were used: an isotype control and SQT01. SQP22 was dosed 4 hours after Fab-Tz infusion. **B-C**, Tumor volumes of NCI-N87 xenografts in SCID mice (**B**) and body weight change (**C**) after dosing with vehicle, distimab vedotin, and SQP22 protodrug alone or with isotype or SQT01. Shown are means ± SEM (*n* = 6 mice/group). *P*-values were determined by two-way ANOVA with Bonferroni correction for multiple comparisons on Day 27. ANOVA, analysis of variance; Fab-Tz, Fab-tetrazine; Isotype Fab-Tz, Isotype Fab-tetrazine; MTD, maximum tolerated dose.

Little to no body weight loss was observed in the group treated with SQP22 and SQT01 compared with the group treated with vehicle control (*P* > 0.99), suggesting minimal non-specific activation of SQP22 (Figure 4C). We also confirmed that treatment with SQT01 alone had no effect on tumor growth or body weight compared with vehicle treatment (Figure S5).

Variables such as dose level, timing, and frequency for SQT01 and SPQ22 are being evaluated to reach an optimal schedule for administration. These results support the conclusion that binding to HER2 by SQT01 effectively localized tetrazine activators at the tumor and led to release of MMAE.

## Conclusion

The data presented in this manuscript together with our previously published work^17^ highlight the power of click chemistry. A single tumor-targeting agent (SQL70 biopolymer) has been shown to activate two different protodrugs: SQP33, with a doxorubicin payload,^17^ and now SQP22, with a MMAE payload. In addition, the results demonstrate how a single protodrug (SQP22) can be activated by two different tumor-targeting agents with distinct dosing methods (intratumoral injection for SQL70 and systemic infusion for SQT01). The biopolymer targeting agent makes the treatment agnostic to tumor type as well as inherent differences in tumor biology that can vary within a single tumor and even more from patient to patient. In other settings, tumor biology can be exploited for specific targeting of the payload, for example, by using SQT01 to target HER2-positive tumors. The treatment of solid tumors remains a significant challenge and the two activating agents represent complementary targeting strategies.

The CAPAC platform is characterized by several features, which unlock unique benefits. First, the use of click chemistry in humans differentiates this approach from biology-based conditional activation strategies. The activation of the protodrug is not dependent on biological factors, but rather is based on the fast, specific, and efficient tetrazine ligation reaction. Nor is the activator required to be taken up by cells allowing non-internalizing or extracellular antigens found in the tumor microenvironment to be used for tumor targeting. In addition, the reliance on chemistry rather than biology is expected to improve the translatability of a therapeutic across animal species and humans, as observed with the lead CAPAC compound SQ3370^17-19^ (and manuscript in preparation).

Second, the targeting agent is decoupled from the payload. This creates a modular system, which enables several beneficial attributes. To start, each component can be optimized for its specific task. The targeting agent can be given in sufficient doses to maximize the probability of saturating the desired antigens in the tumor or tumor microenvironment without the liability of an attached payload. Additionally, the payload can be dosed to maximize its effect (e.g., doses at day 1, 2 or 3) as long as the targeting agent with tetrazine remains at the tumor.

Third, a targeting agent can activate any protodrug that has been previously created. Once a new tumor targeting agent is created, it can be tested in the relevant models with an array of protodrugs, differing in, for example, potency or mechanisms of action, leading to critical insights in choosing a development candidate (Figure 5). Furthermore, the development path is simplified by using off-the shelf protodrugs, which may have already been tested *in vivo* or even in clinical trials. As multiple protodrugs can be used with the same targeting agent, unique combinations and sequencing of therapies are possible that are currently prevented by overlapping toxicities.

**Figure 5.**
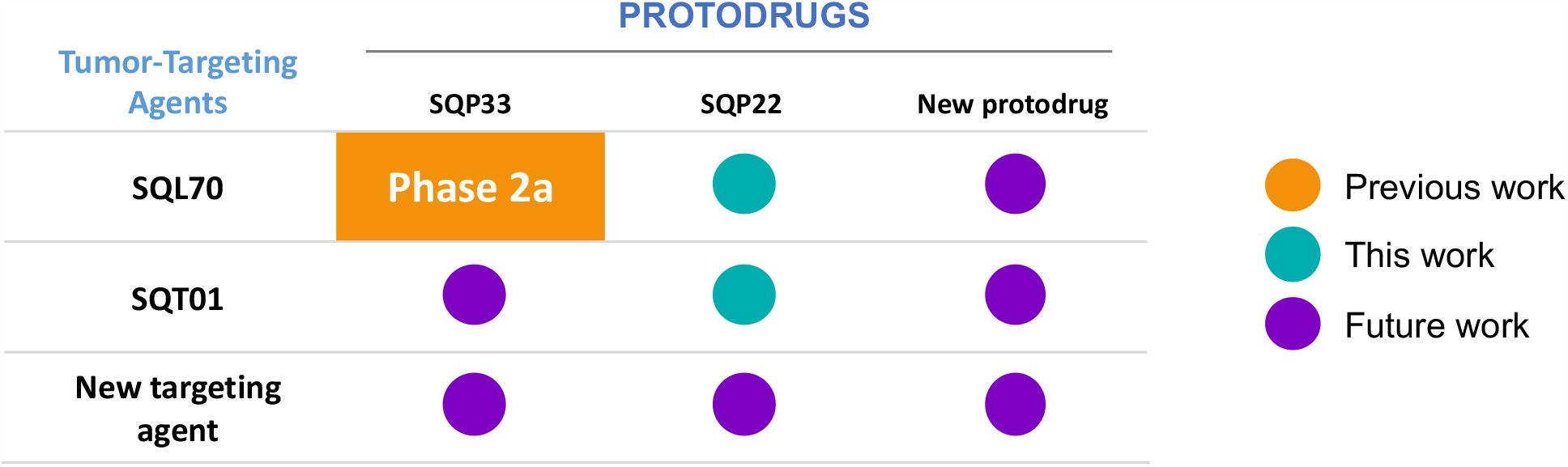
The flexibility of click chemistry for the targeted activation of drugs enables a modular platform in which a single targeting agent can be used with multiple protodrugs or vice versa. Thus, with the addition of each new protodrug or targeting agent, multiple new potential therapeutics are possible, each with the potential benefits unlocked by the platform.

Finally, the drug concentrations at the tumor achieved with our approach can unlock new biological effects. In the phase 1 clinical trial of SQ3370, the treatment has been reported to induce immune activation^19^ that may enhance a systemic anti-tumor response.^17^ Preliminary data with SQP22 and SQL70 have shown similar immune activation effects in the RENCA syngeneic murine model.^41^ Many questions still need to be addressed to further our understanding of the SQP22 protodrug and SQT01, both alone and in combination. Pharmacokinetic and biodistribution studies of SPQ22 and SQT01 will enable optimization of the dosing parameters such as the dose levels, timing, and schedule. Ultimately, the translatability of the safety and efficacy of SQP22 with either SQL70 or SQT01 from animal models to humans will have to be tested.

To summarize, we have demonstrated the modularity and versatility of the CAPAC platform. Anti-tumor activity of the *trans*-cyclooctene-modified-MMAE compound SQP22 in combination with either the intratumorally-injected SQL70 biopolymer or the Fab-directed SQT01 conjugate was observed in multiple preclinical models. These results highlight the power of click chemistry to precisely control the activation of drugs at tumor sites using the tetrazine ligation reaction, as well as support the hypothesis that this approach is agnostic to the format, molecular composition, or delivery method of the targeting agent or payload. Future work on investigational new drug application-enabling studies for SQP22, the development of new targeting agents and payloads, including non-cytotoxic payloads, will further validate the CAPAC platform technology and its potential to engineer new characteristic for therapeutics within biological systems.

## Data Availability

The data generated in this study are available within the article and its Electronic Supporting Information files.

## Authors’ contributions

**Jesse McFarland:** Conceptualization, data curation, investigation, visualization, writing–original draft, writing–review and editing.

**Maša Alečković:** Conceptualization, data curation, formal analysis, investigation, visualization, writing– original draft, writing–review and editing.

**George Coricor:** Conceptualization, data curation, formal analysis, investigation, visualization, writing– original draft, writing–review and editing.

**Sangeetha Srinivasan:** Conceptualization, investigation, visualization, writing–original draft, writing– review and editing.

**Matthew Tso:** Data curation, formal analysis, investigation, visualization, writing–review and editing.

**John Lee:** Writing–review and editing.

**Tri-Hung Nguyen**: Conceptualization, data curation, formal analysis, investigation, visualization, writing–original draft, writing–review, and editing.

**José M. Mejía Oneto:** Conceptualization, funding acquisition, investigation, writing–review and editing.

**Conflicts of Interest**

Jesse M. McFarland, Maša Alečković, George Coricor, Sangeetha Srinivasan, Matthew Tso, John Lee, Tri-Hung Nguyen, and José M. Mejía Oneto are paid employees and shareholders of Shasqi Inc. José M. Mejía Oneto is the Founder and CEO of Shasqi Inc.

## Acknowledgments

This study was funded by Shasqi, Inc. The authors thank Steve Abella, M.D., Scott Wieland, Ph.D., and Sadie Whitaker, Ph.D for critical review of the manuscript. Medical writing support was provided by Martha Mutomba, on behalf of Shasqi, Inc.

## Live subject statement

The described animal studies were performed in accordance with the guidance of the Association for Assessment and Accreditation of Laboratory Animal Care (AAALAC) and approved by the Institutional Animal Care and Use Committee (IACUC) of WuXi AppTec (Nantong, China) or Cephrim Biosciences Inc. (Woburn, MA, USA).

## References

1. J. C. Jewett and C. R. Bertozzi, Cu-free click cycloaddition reactions in chemical biology, Chem Soc Rev, 2010, 39, 1272–1279.

2. The Royal Swedish Academy of Sciences. Click chemistry and bioorthogonal chemisty. The Noble Prize Committee of Chemistry. 5 October 2022. Available at: https://www.nobelprize.org/uploads/2022/10/advanced-chemistryprize2022-2.pdf. Accessed 8 March 2023.

3. K. Wu, N. A. Yee, S. Srinivasan, A. Mahmoodi, M. Zakharian, J. M. Mejia Oneto and M. Royzen, Click activated protodrugs against cancer increase the therapeutic potential of chemotherapy through local capture and activation, Chem Sci, 2021, 12, 1259–1271.

4. M. L. Blackman, M. Royzen and J. M. Fox, Tetrazine ligation: fast bioconjugation based on inverse-electron-demand Diels-Alder reactivity, J Am Chem Soc, 2008, 130, 13518–13519.

5. J. M. Mejía Oneto, M. Gupta, J. K. Leach, M. Lee and J. L. Sutcliffe, Implantable biomaterial based on click chemistry for targeting small molecules, Acta Biomater, 2014, 10, 5099–5105.

6. J. M. Mejia Oneto, I. Khan, L. Seebald and M. Royzen, In vivo bioorthogonal chemistry enables local hydrogel and systemic pro-drug to treat soft tissue sarcoma, ACS Cent Sci, 2016, 2, 476–482.

7. R. Rossin, P. R. Verkerk, S. M. van den Bosch, R. C. Vulders, I. Verel, J. Lub and M. S. Robillard, In vivo chemistry for pretargeted tumor imaging in live mice, Angew Chem Int Ed Engl, 2010, 49, 3375–3378.

8. R. Rossin, S. M. van den Bosch, W. Ten Hoeve, M. Carvelli, R. M. Versteegen, J. Lub and M. S. Robillard, Highly reactive trans-cyclooctene tags with improved stability for Diels-Alder chemistry in living systems, Bioconjug Chem, 2013, 24, 1210–1217.

9. R. Rossin, R. M. Versteegen, J. Wu, A. Khasanov, H. J. Wessels, E. J. Steenbergen, W. Ten Hoeve, H. M. Janssen, A. van Onzen, P. J. Hudson and M. S. Robillard, Chemically triggered drug release from an antibody-drug conjugate leads to potent antitumour activity in mice, Nat Commun, 2018, 9, 1484.

10. N. K. Devaraj, G. M. Thurber, E. J. Keliher, B. Marinelli and R. Weissleder, Reactive polymer enables efficient in vivo bioorthogonal chemistry, Proc Natl Acad Sci U S A, 2012, 109, 4762–4767.

11. N. K. Devaraj, R. Upadhyay, J. B. Haun, S. A. Hilderbrand and R. Weissleder, Fast and sensitive pretargeted labeling of cancer cells through a tetrazine/trans-cyclooctene cycloaddition, Angew Chem Int Ed Engl, 2009, 48, 7013–7016.

12. J. Ferlay, M. Colombet, I. Soerjomataram, D. M. Parkin, M. Piñeros, A. Znaor and F. Bray, Cancer statistics for the year 2020: An overview, Int J Cancer, 2021, DOI: 10.1002/ijc.33588.

13. A. Haslam and V. Prasad, Estimation of the percentage of US patients with cancer who are eligible for and respond to checkpoint inhibitor immunotherapy drugs, JAMA Netw Open, 2019, 2, e192535.

14. P. S. Hegde and D. S. Chen, Top 10 challenges in cancer immunotherapy, Immunity, 2020, 52, 17–35.

15. F. Tian, Y. Lu, A. Manibusan, A. Sellers, H. Tran, Y. Sun, T. Phuong, R. Barnett, B. Hehli, F. Song, M. J. DeGuzman, S. Ensari, J. K. Pinkstaff, L. M. Sullivan, S. L. Biroc, H. Cho, P. G. Schultz, J. DiJoseph, M. Dougher, D. Ma, R. Dushin, M. Leal, L. Tchistiakova, E. Feyfant, H. P. Gerber and P. Sapra, A general approach to site-specific antibody drug conjugates, Proc Natl Acad Sci U S A, 2014, 111, 1766–1771.

16. S. Baah, M. Laws and K. M. Rahman, Antibody-drug conjugates-a tutorial review, Molecules, 2021, 26, 2943.

17. S. Srinivasan, N. A. Yee, K. Wu, M. Zakharian, A. Mahmoodi, M. Royzen and J. M. M. Oneto, SQ3370 activates cytotoxic drug via click chemistry at tumor and elicits sustained responses in injected &non-injected lesions, Adv Ther (Weinh), 2021, 4, 2000243.

18. S. P. Chawla, K. Batty, M. Aleckovic, V. Bhadri, N. Bui, A. D. Guminski, J. M. M. Oneto, S. Srinivasan, J. F. Strauss, V. Subbiah, M. C. Weiss, R. Wilson, N. A. Yee, M. Zakharian and V. Kwatra, Interim phase 1 results for SQ3370 in advanced solid tumors, J Clin Oncol, 2022, 40 (issue 16_suppl), Abstract 3085. DOI: https://ascopubs.org/doi/abs/3010.1200/JCO.2022.3040.3016_suppl.3085. Poster available at:https://www.shasqi.com/assets/pdf/2022-3004-3011-Clinical_Poster_AACR.pdf.

19. S. P. Chawla, K. Batty, M. Alečković, V. A. Bhadri, N. Bui, A. Guminski, J. M. Oneto, S. Srinivasan, J. Strauss, V. Subbiah, M. C. Weiss, R. Wilson, N. Yee, M. Zakharian and V. Kwatra, 1499P Phase I clinical &immunologic data of SQ3370 in advanced solid tumors, Ann Oncol, 2022, 33, Abstract S1232. DOI:1210.1016/j.annonc.2022.1207.1602. Poster available at: https://clin.larvol.com/abstract-detail/ESMO%202022/58818264/58256747.

20. L. Buckel, E. N. Savariar, J. L. Crisp, K. A. Jones, A. M. Hicks, D. J. Scanderbeg, Q. T. Nguyen, J. K. Sicklick, A. M. Lowy, R. Y. Tsien and S. J. Advani, Tumor radiosensitization by monomethyl auristatin E: mechanism of action and targeted delivery, Cancer Res, 2015, 75, 1376–1387.

21. R. Bai, G. R. Pettit and E. Hamel, Dolastatin 10, a powerful cytostatic peptide derived from a marine animal. Inhibition of tubulin polymerization mediated through the vinca alkaloid binding domain, Biochem Pharmacol, 1990, 39, 1941–1949.

22. K. M. Bajjuri, Y. Liu, C. Liu and S. C. Sinha, Tthe legumain protease-activated auristatin prodrugs suppress tumor growth and metastasis without toxicity, ChemMedChem, 2011, 6, 54–59.

23. U. Vaishampayan, M. Glode, W. Du, A. Kraft, G. Hudes, J. Wright and M. Hussain, Phase II study of dolastatin-10 in patients with hormone-refractory metastatic prostate adenocarcinoma, Clin Cancer Res, 2000, 6, 4205–4208.

24. Y. Liu, K. M. Bajjuri, C. Liu and S. C. Sinha, Targeting cell surface alpha(v)beta(3) integrin increases therapeutic efficacies of a legumain protease-activated auristatin prodrug, Mol Pharm, 2012, 9, 168–175.

25. H. Li, C. Yu, J. Jiang, C. Huang, X. Yao, Q. Xu, F. Yu, L. Lou and J. Fang, An anti-HER2 antibody conjugated with monomethyl auristatin E is highly effective in HER2-positive human gastric cancer, Cancer Biol Ther, 2016, 17, 346–354.

26. J. A. Francisco, C. G. Cerveny, D. L. Meyer, B. J. Mixan, K. Klussman, D. F. Chace, S. X. Rejniak, K. A. Gordon, R. DeBlanc, B. E. Toki, C. L. Law, S. O. Doronina, C. B. Siegall, P. D. Senter and A. F. Wahl, cAC10-vcMMAE, an anti-CD30-monomethyl auristatin E conjugate with potent and selective antitumor activity, Blood, 2003, 102, 1458–1465.

27. ADCETRIS (brentuximab vedotin) prescribing information (2011) Seagen Inc., Bothwell, WA, USA. Available at: https://www.accessdata.fda.gov/drugsatfda_docs/label/2014/125388_s056s078lbl.pdf. Accessed 8 March 2023.

28. PADCEV (enfortumab vedotin-ejfv) prescribing information (2019) Seagen Inc., Bothell, WA, USA. Available at: https://www.accessdata.fda.gov/drugsatfda_docs/label/2019/761137s000lbl.pdf. Accessed 8 March 2023.

29. POLIVY (polatuzumab vedotin-piiq) prescribing information (2019) Genentech, South San Francisco, CA, USA. Available at: https://www.accessdata.fda.gov/drugsatfda_docs/label/2019/761137s000lbl.pdf. Accessed 8 March 2023.

30. TIVDAK (tisotumab vedotin-tftv) prescribing information (2021) Seagen Inc., Bothell, WA, USA. Available at: https://www.accessdata.fda.gov/drugsatfda_docs/label/2021/761208Orig1s000lbledt.pdf. Accessed 8 March 2023).

31. P. Schöffski, J. P. Delord, E. Brain, J. Robert, H. Dumez, J. Gasmi and A. Trouet, First-in-man phase I study assessing the safety and pharmacokinetics of a 1-hour intravenous infusion of the doxorubicin prodrug DTS-201 every 3 weeks in patients with advanced or metastatic solid tumours, Eur J Cancer, 2017, 86, 240–247.

32. M. Poreba, Protease-activated prodrugs: strategies, challenges, and future directions, FEBS J, 2020, 287, 1936–1969.

33. M. M. Mita, R. B. Natale, E. M. Wolin, B. Laabs, H. Dinh, S. Wieland, D. J. Levitt and A. C. Mita, Pharmacokinetic study of aldoxorubicin in patients with solid tumors, Invest New Drugs, 2015, 33, 341–348.

34. Y. Pei, M. Li, Y. Hou, Y. Hu, G. Chu, L. Dai, K. Li, Y. Xing, B. Tao, Y. Yu, C. Xue, Y. He, Z. Luo and K. Cai, An autonomous tumor-targeted nanoprodrug for reactive oxygen species-activatable dual-cytochrome c/doxorubicin antitumor therapy, Nanoscale, 2018, 10, 11418–11429.

35. W. D. Tap, Z. Papai, B. A. Van Tine, S. Attia, K. N. Ganjoo, R. L. Jones, S. Schuetze, D. Reed, S. P. Chawla, R. F. Riedel, A. Krarup-Hansen, M. Toulmonde, I. Ray-Coquard, P. Hohenberger, G. Grignani, L. D. Cranmer, S. Okuno, M. Agulnik, W. Read, C. W. Ryan, T. Alcindor, X. F. G. Del Muro, G. T. Budd, H. Tawbi, T. Pearce, S. Kroll, D. K. Reinke and P. Schöffski, Doxorubicin plus evofosfamide versus doxorubicin alone in locally advanced, unresectable or metastatic soft-tissue sarcoma (TH CR-406/SARC021): an international, multicentre, open-label, randomised phase 3 trial, Lancet Oncol, 2017, 18, 1089–1103.

36. Charles River research models (technical sheet). NOD SCID mouse hematology (2022) Charles River Inc., Wilmington, MA, USA. Available at: https://www.criver.com/sites/default/files/resources/NODSCIDMouseClinicalPathologyData.pdf. Accessed 8 March 2023.

37. F. Shi, Y. Liu, X. Zhou, P. Shen, R. Xue and M. Zhang, Disitamab vedotin: a novel antibodydrug conjugates for cancer therapy, Drug Deliv, 2022, 29, 1335–1344.

38. Y. Hu, Y. Zhu, X. Wei, C. Tang and W. Zhang, Disitamab vedotin, a novel HER2-directed antibody-drug conjugate in gastric cancer and other solid tumors, Drugs Today (Barc), 2022, 58, 491–507.

39. Z. Peng, T. Liu, J. Wei, A. Wang, Y. He, L. Yang, X. Zhang, N. Fan, S. Luo, Z. Li, K. Gu, J. Lu, J. Xu, Q. Fan, R. Xu, L. Zhang, E. Li, Y. Sun, G. Yu, C. Bai, Y. Liu, J. Zeng, J. Ying, X. Liang, N. Xu, C. Gao, Y. Shu, D. Ma, G. Dai, S. Li, T. Deng, Y. Cui, J. Fang, Y. Ba and L. Shen, Efficacy and safety of a novel anti-HER2 therapeutic antibody RC48 in patients with HER2-overexpressing, locally advanced or metastatic gastric or gastroesophageal junction cancer: a single-arm phase II study, Cancer Commun (Lond), 2021, 41, 1173–1182.

40. C. M. McKertish and V. Kayser, Advances and limitations of antibody drug conjugates for cancer, Biomedicines, 2021, 9.

41. Aleckovic M, Srinivasan S, McFarland J, et al954 SQ2270, a novel MMAE-based therapeutic, promotes tumor growth inhibition and extensive immune cell infiltration in the RENCA cancer model. J ImmunoTherapy of Cancer 2022;10:doi: 10.1136/jitc-2022-SITC2022.0954. Poster available at: https://shasqi.com/assets/pdf/2022-11-07_-_SITC_SQ2270_Poster.pdf. Accessed 8 March 2023.

