## Supplemental Information for "Click chemistry selectively activates an auristatin protodrug with either intratumoral or systemic tumor-targeting agents"


| **Table of Contents** | **Page number** |
|---|---|
| Materials and Methods | S2-S12 |
| **Table S1.** SQP22 stability in human and mouse plasma | S13 |
| **Figure S1.** SQP22 protodrug stability in tissue homogenates | S14 |
| **Figure S2.** SQP22 with SQL70 leads to complete regression of Karpas 299 tumors | S15 |
| **Figure S3.** SQP22 with SQL70 leads to reduced body weight loss compared to MMAE | S16 |
| **Figure S4.** Characterization of SQT01 | S17 |
| **Figure S5.** SQT01 has no effect on tumor volume or mouse body weight | S18 |

### Materials and methods

**Safety statement**: no unexpected or unusually high safety hazards were encountered in the execution of the experiments for this manuscript.

#### *Preparation of SQP22 protodrug, SQL70 biopolymer, and the HER2 Fab-tetrazine conjugate SQT01*

*Synthesis of SQP22*

The synthesis of the SQP22 protodrug is described in detail below (Supplementary Scheme 1).

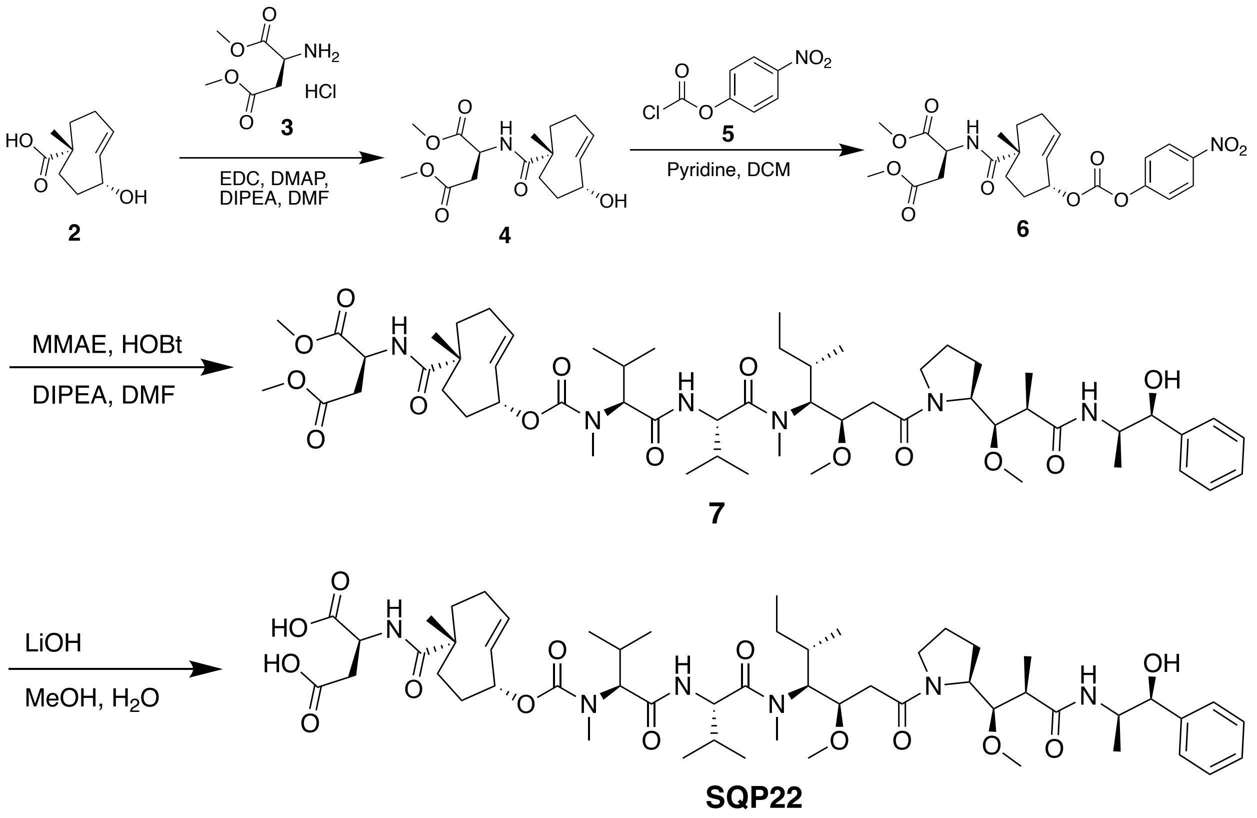


**Scheme 1.** Synthesis of SQP22 protodrug.

*Preparation of compound 4.* The steps for SQP22 synthesis are shown in Scheme 1. Compound **2** is synthesized using established methods.[1] To a solution of compound **2** (1.00 g, 5.43 mmol) in DCM (10 mL) was added DIPEA (2.10 g, 16.3 mmol), EDCI (2.08 g, 10.9 mmol) and DMAP (1.33 g, 10.9 mmol) and compound **3** (1.61 g, 8.14 mmol). The mixture was stirred at 25 °C for 16 hrs. TLC indicated compound **2** was consumed completely and one new spot formed. The reaction mixture was partitioned between DCM

(20 mL) and $H_2O$ (10 mL). The organic phase was separated, washed with saturated aqueous citric acid (3 mL) and brine (20 mL), then dried over $Na_2SO_4$, filtered, and concentrated under reduced pressure to give a residue. The residue was purified by column chromatography (Petroleum ether/Ethyl acetate = 3/1 to 1/1) to give compound **4** (700 mg, 39.4% yield) as a white oil.

**$^1$HNMR** (400MHz, $CDCl_3$):

δ ppm 1.12 (s, 3 H), 1.60 (dd, *J* =15.45, 6.19 Hz, 1 H), 1.79 - 1.87 (m, 2 H), 1.92 (br d, *J* = 5.88 Hz, 1 H), 1.95 (s, 1 H), 1.98 - 2.00 (m, 1 H), 2.02 (br d, *J* = 4.13 Hz, 1 H), 2.26 (dd, *J* = 11.63, 3.88 Hz, 1 H), 2.30 - 2.36 (m, 1 H), 2.77 - 2.89 (m, 1 H), 2.88 - 2.88 (m, 1 H), 3.00 (dd, *J* = 16.95, 4.57 Hz, 1 H), 3.70 (s, 4 H), 3.75 (s, 3 H), 4.80 (dt, *J* = 8.00, 4.50 Hz, 1 H), 5.66 (dd, *J* = 16.63, 2.38 Hz, 1 H), 6.02 - 6.12 (m, 1 H), 6.54 (br d, *J* = 7.88 Hz, 2 H).

*Preparation of Compound 6:* To a solution of compound **4** (700 mg, 2.14 mmol) in DCM (5 mL) was added pyridine (846 mg, 10.7 mmol) and compound **5** (1.72 g, 8.55 mmol) in DCM (5 mL). The mixture was stirred at 25 °C for 1 hrs. TLC indicated compound **4** was consumed completely and one new spot formed. The reaction mixture was partitioned between DCM (20 mL) and $H_2O$ (10 mL). The organic phase was separated, washed with saturated aqueous citric acid (3 mL) and brine (20 mL), dried over $Na_2SO_4$, filtered, and concentrated under reduced pressure to give a residue. The residue was purified by column chromatography (Petroleum ether/Ethyl acetate = 3/1 to 1/1) to give compound **6** (490 mg, 46.5% yield) as a yellow oil.

**$^1$HNMR** (400MHz, $CDCl_3$):

δ ppm 1.17 (s, 3 H), 1.55 - 1.61 (m, 1 H), 1.58 (br s, 1 H), 1.76 (dd, *J* = 14.76, 6.25 Hz, 1 H), 1.87 - 2.03 (m, 3 H), 2.06 - 2.15 (m, 1 H), 2.19 - 2.41 (m, 3 H), 2.82 (dd, *J* = 17.13, 4.50 Hz, 1 H), 3.03 (dd, *J* = 17.07, 4.44 Hz, 1 H), 3.72 (s, 3 H), 3.77 (s, 3 H), 4.78 - 4.86 (m, 1 H), 5.67 (dd, *J* = 16.70, 2.44 Hz, 1 H), 6.03 - 6.14 (m, 1 H), 6.58 (br d, *J* = 7.88 Hz, 1 H), 7.40 - 7.45 (m, 2 H), 8.27 - 8.33 (m, 2 H).

*Preparation of compound 7:* To a solution of compound **6** (490 mg, 995 µmol) and MMAE (714 mg, 995 µmol) in DMF (4 mL) was added DIPEA (64.3 mg, 497 µmol) and HOBt (202 mg, 1.49 mmol). The mixture was stirred at 25 °C for 16 hrs. LC-MS showed that compound **6** was consumed completely, and one main peak with desired mass was detected. The residue was purified by prep-HPLC (0.1% TFA conditions) to give compound **7** (500 mg, 46.9% yield) as a white solid.

**$^1$HNMR** (400MHz, $CDCl_3$):

δ ppm 0.84 (br d, *J* = 6.75 Hz, 4 H), 0.89 (br d, *J* = 4.50 Hz, 5 H), 0.92 (br d, *J* = 6.63 Hz, 4 H), 0.98 (br d, *J* = 6.25 Hz, 3 H), 1.04 (br d, *J* = 6.88 Hz, 3 H), 1.16 (s, 3 H), 1.25 - 1.27 (m, 3 H), 1.59 - 1.74 (m, 3 H), 1.88 (br d, *J* = 9.38 Hz, 4 H), 2.07 (br d, *J* = 8.38 Hz, 5 H), 2.27 (br s, 4 H), 2.36 - 2.43 (m, 2 H), 2.45 - 2.53 (m, 1 H), 2.89 (br s, 6 H), 2.95 - 3.01 (m, 4 H), 3.04 (br s, 2 H), 3.29 - 3.34 (m, 3 H), 3.36 - 3.47 (m, 5 H), 3.67 - 3.73 (m, 4 H), 3.76 (s, 3 H), 3.82 - 3.89 (m, 1 H), 4.05 - 4.19 (m, 3 H), 4.28 (br s, 1 H), 4.63 - 4.86 (m, 3 H), 4.96 (d, *J* = 2.50 Hz, 1 H), 5.24 (br s, 1 H), 5.63 (br d, *J* = 18.14 Hz, 1 H), 5.82 (br s, 1 H), 6.53 - 6.74 (m, 3 H), 7.30 - 7.41 (m, 5 H).

*Preparation of SQP22*: To a solution of compound **7** (500 mg, 0.47 mmol) in MeOH (5 mL) was added $LiOH \cdot H_2O$ (196 mg, 4.67 mmol) in $H_2O$ (2 mL). The mixture was stirred at 25 °C for 16 hrs. LC-MS showed compound **7** was consumed completely, and one main peak with desired mass was detected. The residue was adjusted pH ~ 2 with saturated aqueous citric acid, then purified by prep-HPLC (0.1% TFA condition) to give **SQP22** (265 mg 53.4% yield) as a white solid.

**$^1$HNMR** (400MHz, $CDCl_3$):

δ ppm 0.80 - 1.04 (m, 25 H), 1.09 (s, 3 H), 1.24 (d, *J* = 6.88 Hz, 3 H), 1.69 - 1.82 (m, 2 H), 1.88 - 1.94 (m, 3 H), 2.02 - 2.11 (m, 4 H), 2.16 (br d, *J* = 18.64 Hz, 1 H), 2.11 - 2.24 (m, 2 H), 2.32 (br d, *J* = 5.25 Hz, 2 H), 2.40 - 2.45 (m, 1 H), 2.52 (br d, *J* = 5.50 Hz, 2 H), 2.81 (br dd, *J* = 14.01, 4.88 Hz, 1 H), 2.94 - 3.04 (m, 2 H), 3.08 (s, 2 H), 3.14 - 3.27 (m, 6 H), 3.33 (s, 1 H), 3.39 (s, 3 H), 3.48 - 3.57 (m, 2 H), 3.94 (br d, *J* = 1.25 Hz, 1 H), 4.05 - 4.18 (m, 4 H), 4.30 (br dd, *J* = 6.19, 4.82 Hz, 2 H), 4.54 - 4.67 (m, 4 H), 4.91 (br d, *J* = 2.00 Hz, 2 H), 5.30 (br s, 1 H), 5.64 - 5.73 (m, 1 H), 5.79 - 5.89 (m, 1 H), 6.61 (br d, *J* = 7.38 Hz, 1 H), 7.30 - 7.42 (m, 5 H), 7.55 - 7.64 (m, 1 H).

*Synthesis of SQL70*

The synthesis of the SQL70 biopolymer has been described previously.[2]

*Generation of the HER2 Fab-tetrazine conjugate* SQT01

*Generation of SQT01*: Coding sequences of the variable region of heavy chain and light chain of trastuzumab (human anti-HER2) antibody were used to generate HER2 Fab-expressing constructs. Coding sequences were synthesized and subcloned into the expression vector. The constructed plasmids were transformed to *E. coli* for propagation and scale-up. Purified plasmids were confirmed by sequencing. The constructs containing the heavy chain and light chain of the HER2 Fab were transfected into HEK293 cells with polymer polyethylenimine (PEI) reagent. The culture medium was harvested at 6-7 days post-transfection. The culture medium containing HER2 Fab was centrifuged, filtered, and then loaded onto KappaSelect affinity column (Mabselect Prism). The loading buffer was 25 mM Tris containing 150 mM NaCl, pH 8.0, and eluted with 100 mM sodium-citrate buffer containing 150 mM NaCl, pH 2.5. The collected solution was neutralized with 1 M arginine, 400 mM succinic acid buffer, pH 9.0. The affinity-purified protein was further purified by gel filtration with Superdex S-200 5/150GL column chromatography. The sample injection was 20 mL with a flow rate of 0.3 mL/min and mobile phase 2X PBS at pH 7.4. The purified HER2 Fab was analyzed by SDS-PAGE and SEC-HPLC.

*Tetrazine conjugation of HER2 Fab:* HER2 Fab was buffer exchanged to PBS pH 7.4 overnight. The methyltetrazine-PEG9-NHS (SiChem #SC-8808) was dissolved in DMSO to make a 10 mM stock solution. For conjugation, the two components were reacted at 3:1 (methyltetrazine-PEG9-NHS to HER2 Fab) molar ratio at 25 °C for 2 hours. Then the solution was buffer exchanged to PBS pH 7.4 to remove excess methyltetrazine-PEG9-NHS and buffer salts.

*Characterization of SQT01:* The sample of prepared SQT01 conjugate was analyzed by SDS-PAGE and LC-MS to confirm the formation of SQT01 conjugate and to determine the tetrazine-to-antibody ratio (Supplementary Fig. 4). Mass spectra and HPLC-SEC traces were obtained for SQT01 and showed > 99% monomeric species for the Fab-tetrazine conjugate with minimal aggregation (≤ 0.3%). The tetrazine-to-antibody ratio was calculated to be 2.2. Cell binding analysis by flow cytometry using NCI-N87 (HER2 positive) cells was tested with either unconjugated HER2 Fab, SQT01, or isotype control. Mean fluorescent intensity (MFI) was calculated and plotted against antibody concentration. Comparable binding was observed with SQT01 compared to unconjugated HER2 Fab.

*Flow cytometry analysis of SQT01:* NCI-N87 (HER2-positive) human gastric cancer cells were collected by centrifugation and resuspended with FACS buffer (PBS containing 2% FBS, pH 7.4). The cells were seeded in a 96-well plate (200,000 cells, 100 μL), the plate was centrifuged at 400 x g for 5 minutes, the supernatant was removed, and the cells were incubated at 4 °C for 1 hour with titration of 0.00002 - 20 mg/mL SQT01, HER2 Fab unconjugated, or isotype control (IgG) solution. The plate was centrifuged, and the cells were washed 3 times with FACS buffer. Then the cells were resuspended in 100 mL solution containing anti-human IgG-Alexa fluor 488 (catalog #11013, Life Technologies, Carlsbad, CA, USA), and incubated in the dark at 4 °C for 1 hour. The supernatant was removed, the cells were washed twice with PBS, and then the cells were analyzed by CytoFLEX flow cytometer (Beckman Coulter, Brea, CA, USA).

***In vitro assays to evaluate SQP22 cytotoxicity and stability in plasma and tissue homogenates***

*Cytotoxicity assay*

The *in vitro* cytotoxicity of SQP22 in combination with methyltetrazine (Click Chemistry Tools, Scottsdale, AZ, USA, #1125) was tested in cancer cell lines (MC38, B6-F10, EMT6, RENCA) using a CellTiter-Glo (CTG) assay (Promega, Madison, WI, USA). MC38 and B16-F10 cells were cultured DMEM with 10% fetal bovine serum (FBS) and 1x penicillin-streptomycin (Pen/Strep), while EMT6 and RENCA were cultured RPMI 1640 supplemented with 10% FBS and 1x Pen/Strep. All cell lines were maintained in an incubator at 37 °C in an atmosphere of 5% $CO_2$. Cells at ~ 60-80% confluency were collected via trypsinization. One thousand MC38, B16-F10, EMT6 cells or two thousand RENCA cells in 95 μL of media were seeded into 96-well plates overnight. 10 mM SQP22 in DMSO was mixed with an equal volume of 10 mM tetrazine in DMSO or DMSO alone to generate 5 mM drug solutions of SQP22 or SQP22/tetrazine. These solutions were aged at ambient temperature for 15 minutes, protected from light, before being further diluted with complete media. Cells were treated at 8 concentrations generated via 5-fold serial dilutions (0.5 μM top concentration) and a non-treatment control. Cells were treated in triplicates for 72 hours before CTG assays.

The CTG assay was performed with the manufacturer's recommended procedure with modifications. Prior to the CTG assay, the plate and CTG reagent were at room temperature. CTG reagent (100 μL) was added to each well of the 96-well plate, and the contents were mixed to induce cell lysis. The plate's luminescent signal was stabilized for 10 minutes before reading by a microplate reader (FlexStation 3, Molecular Device, San Jose, CA, USA), and the signals were used to generate a viability curve of drug concentrations versus cell response. GraphPad Prism software was used to determine the $IC_{50}$ of the parameters on the tested cells.

*Plasma stability assay*

The metabolic stability of SQP22 was assessed in human and mouse plasma. The human and mouse plasma in K2 EDTA were obtained from BioIVT (Westbury, NY, USA). The assay was carried out in 96-well microtiter plates. Compounds were incubated in duplicate at 37 °C in the presence of plasma. Reaction mixtures (50 μL) contained a final concentration of 1 μM test SQP22. The extent of metabolism was calculated as the disappearance of the test compound, compared to the 0-minute control reaction incubations. Propantheline was included as a positive control to verify assay performance.

At each of the four time points, 300 μL of quench solution (50% acetonitrile, 50% methanol, and 0.05% formic acid, warmed up at 37 °C) containing internal standards was added to each well. Plates were sealed, vortexed, and centrifuged at 4 °C for 15 minutes at 4,000 rpm. The supernatant was transferred to fresh plates for LC-MS/MS analysis.

All samples were analyzed by LC-MS/MS using an AB Sciex API 4000 instrument coupled to a Shimadzu LC-20AD LC Pump system. Analytical samples were separated using a Waters Atlantis T3 dC18 reverse phase HPLC column (10 mm x 2.1 mm) at a flow rate of 0.5 mL/min. The mobile phase consisted of 0.1% formic acid in water (solvent A) and 0.1% formic acid in 100% acetonitrile (solvent B).

The extent of metabolism was calculated as the disappearance of the test compound, compared to the 0-minute control reaction incubations. Initial rates were calculated for the compound concentration and used to determine t1/2 values.

*Tissue homogenate stability assay*

For sample preparation and processing, mouse naïve plasma, liver, spleen, heart, and kidney samples were obtained from BioIVT (Westbury, NY, USA). Two volumes of each tissue type in one volume of PBS (weight/volume) were homogenized by bead milling to prepare the homogenates. The samples were then spiked with SQP22 to a concentration of 1 µM and incubated at 37 °C. In duplicate, a 40 µL aliquot was collected at 0, 5, 15, 30 and 60 minutes, and at 2, 4, 8, and 24 hours. The samples, along with calibration samples with 0.5 – 1000 ng/mL of MMAE or SQP22, were processed by adding 4 volumes of ice-cold acetonitrile containing 100 ng/mL MMAF, and then centrifuged at 6,100 x g for 30 minutes. 170 µL of each supernatant was transferred to an autosampler plate insert and dried completely for about 2 hours on medium heat in the dryvac. The samples were then reconstituted with 20 µL of 2 mM ammonium acetate and 0.2% formic acid in water.

Sample analysis was performed using LC-MS/MS system, in which a Shimadzu VP Series HPLC system was in tandem with a SCIEX API 6500 (Foster City, CA, USA). The HPLC was equipped with a 20 x 2 mm Proto 200 C18 column (Higgins Analytical, Mountain View, CA, USA). An injection volume of 12.0 µL was used with a flow rate of 1.2 mL/min, and a gradient mobile phase A (2 mM ammonium acetate in water with 0.2% formic acid) and phase B (0.2% formic acid in acetonitrile). For detecting MMAE and SQP22 by mass spectrometry, a TurboIonSpray (ESI) in positive ionization mode was used with the cone voltage of 450 °C. Transitions monitored were used as follows: MMAE 718.6 –> 686.8 m/z; SQP22: 1043.6 –> 718.5 m/z; MMAF (as internal standard) 732.6 –> 170.3 m/z.

***Animal studies***

All procedures related to animal handling, care, and treatment in the study were performed according to the guidance of the Association for Assessment and Accreditation of Laboratory Animal Care (AAALAC) and approved by the Institutional Animal Care and Use Committee (IACUC) of WuXi AppTec (Nantong, China) or Cephrim Biosciences, Inc.

*Karpas 299 xenograft model*

The murine Karpas 299 (human non-Hodgkin′s large cell lymphoma) xenograft studies were conducted at WuXi AppTec. Karpas 299 cells were expanded in complete culture medium RPMI-1640 containing 20% FBS, 2 mM glutamine, and 1% Pen-Strep in a 37 °C incubator in an atmosphere of 5% $CO_2$. Cells were routinely passaged twice a week. Once the Karpas 299 cells reached the exponential growth phase, cells were harvested, washed, and counted for cell inoculation in mice. Female C.B-17 SCID mice 6-8 weeks old were inoculated subcutaneously at the right upper flank with 1 x $10^6$ Karpas 299 cells in 0.2 mL of PBS with 50% Matrigel (1:1) for tumor development. Animals were distributed into groups of 8 for the efficacy study, and tumors were left to grow to group mean tumor volumes of ~100 $mm^3$.

To determine the attenuation of SQP22 *in vivo*, each group of 8 animals was administered vehicle control (10% hydroxypropyl-beta-cyclodextran [HPCD]) dosed IV on days 1–5; single IV dose of MMAE at 0.5 mg/kg (1x) on day 1; or SQP22 dosed at 7.3 mg/kg on days 1–5 (50-times [50x] molar equivalence of MMAE per dosing cycle) (Fig. 2). To determine activation of SQP22 by the SQL70 biopolymer, each group of 8 animals was administered the vehicle control dosed IV on days 1–5; a single IV dose of MMAE at 0.5 mg/kg (1x) on day 1; SQP22 protodrug dosed at 2.2 mg/kg IV on days 1–5; a single dose of 100 µL of SQL70 biopolymer administered intratumorally followed by SQP22 protodrug dosed at 1.5 or 2.2 mg/kg IV 1 hour later on day 1 then dosed again on days 2, 3, 4, and 5 at 1.5 or 2.2 mg/kg (for 10x or 15x molar equivalents to MMAE [0.5 mg/kg] per dosing cycle of days 1–5) (Fig. 2).

Tumor volume and body weight were measured two times weekly for both experiments. Tumor volume was measured using a caliper and calculated using the following formula: tumor volume ($mm^3$) = length x width$^2$ x 0.5. Complete response to treatment was defined as no palpable tumors measured over 3 consecutive days. Body weight loss was calculated according to the formula: percent body weight loss = (starting weight minus current weight)/(starting weight) x 100. Animals were euthanized when tumors reached 2,000 $mm^3$ or body weight loss > 20% from the initial weight.

*RENCA syngeneic model*

The mouse RENCA (renal adenocarcinoma) syngeneic tumor studies were conducted at Cephrim Biosciences, Inc. RENCA cells were expanded in complete culture medium RPMI-1640 containing 10% FBS, 2 mM glutamine, and 1% Pen-Strep in a 37 °C incubator with 5% $CO_2$. Once RENCA cells reached the exponential growth phase, cells were harvested, washed, and counted for inoculation in mice. Female BALB/c mice 6-8 weeks old were injected subcutaneously at the right flank with 5 x$10^5$ cells in 100 µL of PBS. Animals were divided into groups of 3, 4 or 5 animals for the efficacy study, and tumors were left to grow to group mean tumor volumes of ~90 $mm^3$. The groups were administered vehicle control (10% HPCD) dosed IV on days 1–3, a single intratumoral dose of 40 µL of SQL70 biopolymer on day 1, a single IV dose of MMAE at 1 mg/kg (1x) on day 1, or SQL70 biopolymer dosed intratumorally on day 1 followed by IV dose of SQP22 one hour later at either 2.9 mg/kg or 4.4 mg/kg on days 1–3 (i.e., 6x, or 9x molar equivalents to MMAE [1 mg/kg] per dosing cycle, respectively) (Fig. 3).

Tumor volume and body weight were measured three times weekly, with tumor volume and body weight change calculated as described for the Karpas 299 xenograft model above. Animals were euthanized when tumors reached 2,000 $mm^3$ or significant body weight loss of > 20% from the initial weight. For complete blood count, ~20 µL of whole blood was collected in EDTA tubes. Complete blood count was performed using the Drew Scientific HEMAVET Multispecies Hematology Analyzer according to the manufacturer's instruction (Delta Scientific Inc., Miami Lakes, FL, USA).

*NCI-N87 xenograft model*

The NCI-N87 (gastric carcinoma) xenograft studies were conducted at Shanghai ChemPartner (Shanghai, China). NCI-N87 cells were cultured in RPMI-1640 supplemented with 10% FBS and 1% Pen-Strep in a

37 °C incubator with 5% $CO_2$. Cells were harvested and counted for tumor inoculation when the cells reached the exponential growth phase. Female C.B-17 SCID mice 6-8 weeks old were inoculated subcutaneously at the right flank with 3 x $10^6$ cells in a 200 µL mixture of RPMI-1640 with Matrigel (1:1) for tumor development. Animals were divided into groups of 6 for the efficacy study, and tumors were left to grow to tumor volume of ~100-150 $mm^3$ prior to dosing. All doses were given IV. Each group of 6 animals was administered the vehicle control (10% HPCD) on days 1–3; SQP22 alone at 5 mg/kg on days 1–3; a single dose of isotype Fab-tetrazine at 50 mg/kg on day 1 followed by SQP22 at 5 mg/kg on day 1 (4 hours after isotype conjugate) as well as on days 2 and 3; or a single dose of SQT01 at 50 mg/kg on day 1 followed by SQP22 at 5 mg/kg on day 1 (4 hours after the SQT01), as well as on days 2 and 3 (Fig. 4). For a positive control group, animals received a single dose of disitamab-vedotin (Catalog # HY-P9985, MedChemExpress, Monmouth Junction, NJ, USA) at 10 mg/kg on day 1 (Fig. 4).

Tumor volume was measured three times weekly in two dimensions using a caliper. Body weight was collected daily for the first week and twice weekly after the first week. Tumor volume and body weight change were calculated as previously described for the Karpas 299 xenograft model. Animals were euthanized when tumors reached 2,000 $mm^3$ or significant body weight loss of > 20% from the initial weight.

#### *Statistical Analysis*

Continuous variables are expressed as mean ± standard error of the mean (SEM), unless otherwise noted. For tumor volume and body weight changes, data were analyzed by two-way analysis of variance (ANOVA) with Bonferroni correction for multiple comparisons. For complete blood counts and maximum acute body weight change, data were analyzed using an ordinary one-way ANOVA with Bonferroni correction for multiple comparisons. In all cases, significance was defined as $P \leq 0.05$. Statistical analysis was carried out using GraphPad Prism 9 Software.

**References**

1. Wu, N. A. Yee, S. Srinivasan, A. Mahmoodi, M. Zakharian, J. M. Mejia Oneto and M. Royzen, Click activated protodrugs against cancer increase the therapeutic potential of chemotherapy through local capture and activation, *Chem Sci*, 2021, 12, 1259-1271.

2. S. Srinivasan, N. A. Yee, K. Wu, M. Zakharian, A. Mahmoodi, M. Royzen and J. M. M. Oneto, SQ3370 activates cytotoxic drug via click chemistry at tumor and elicits sustained responses in injected & non-injected lesions, *Adv Ther (Weinh)*, 2021, 4, 2000243.

### Supplementary results

**Table S1.** SQP22 stability in human and mouse plasma

| Plasma Type | SQP22 Remaining After 4 Hour Incubation (%) |
|---|---|
| Human | 94 |
| Mouse | 100 |

SQP22 protodrug was incubated in human or mouse plasma for 4 hours at 37 °C and the fraction remaining was quantified by LC-MS. LC-MS, liquid chromatography-mass spectrometry.

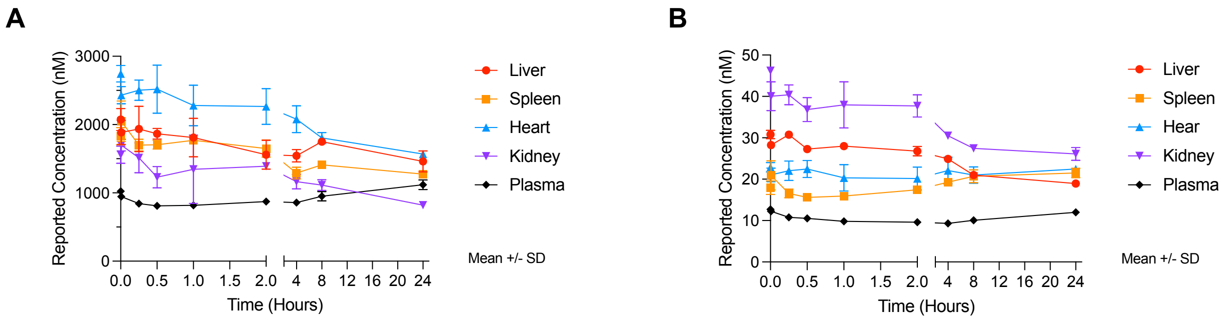


**<u>Figure S1.</u>** SQP22 protodrug stability in tissue homogenates. **A-B**, SQP22 was incubated at 37 °C in various tissue homogenates; samples were taken at the indicated time points and then analyzed for the concentration of SQP22 **(A)** and MMAE (**B**). Differences in apparent SQP22 starting concentrations are due to quantification with a standard curve prepared in plasma, rather than the matched tissue homogenate. MMAE present at time 0 represents a small amount of contaminant resulting from the synthesis of SQP22. Values shown are mean of replicates ± SD (*n* = 2 replicates/time point). MMAE, monomethyl auristatin E; SD, standard error.

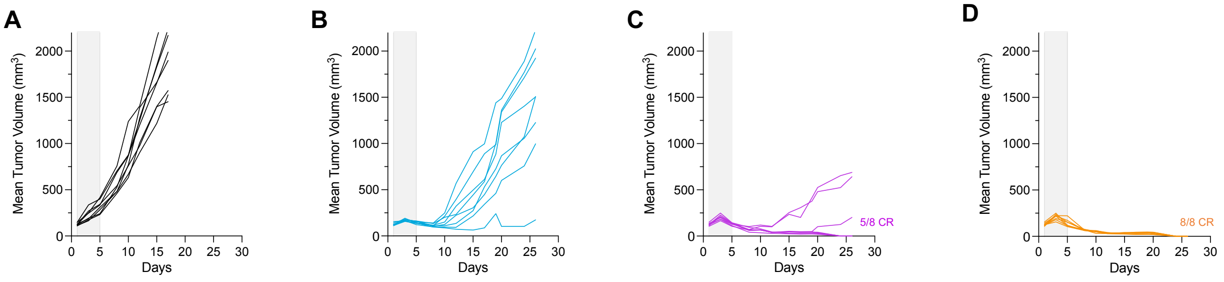


**Figure S2.** SQP22 with SQL70 leads to complete regression of Karpas 299 tumors. **A-D**, Individual tumor volumes in Karpas 299-bearing animals treated with vehicle (**A**), 0.5 mg/kg MMAE (**B**) or SQP22 at a cumulative dose of 10x (**C**) or 15x (**D**) molar equivalents of MMAE based on the dosing schedule described in Fig. 2D.

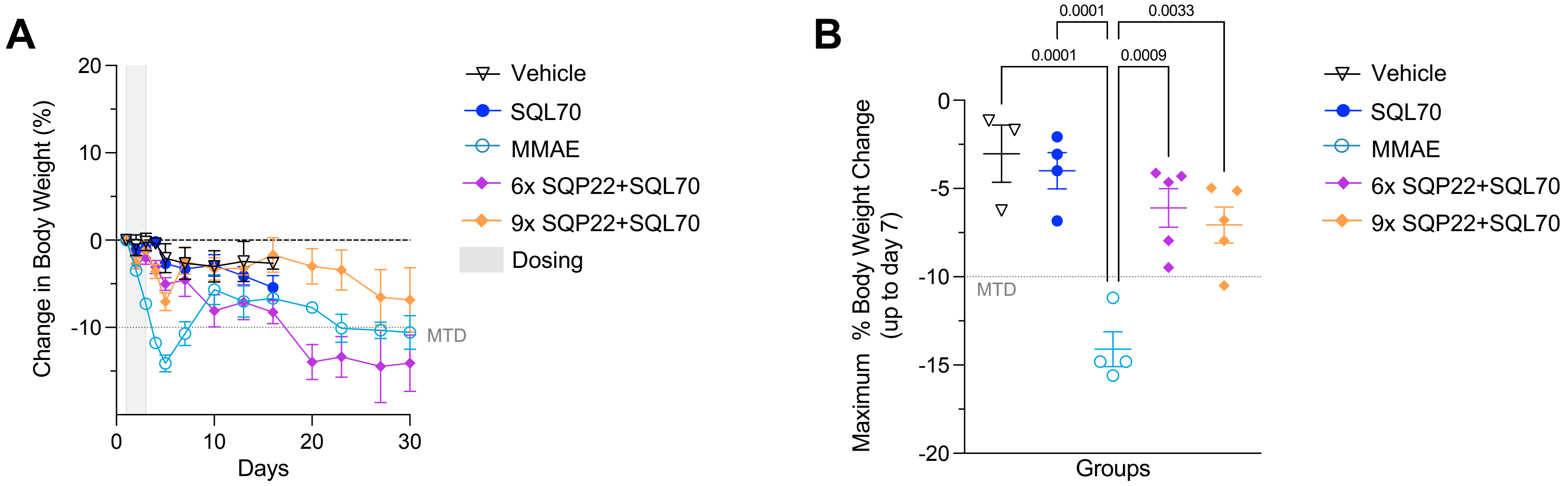


**<u>Figure S3.</u>** SQP22 with SQL70 leads to reduced body weight loss compared to MMAE. **A**, Body weight changes of BALB/c mice bearing RENCA tumors following treatment with vehicle (*n* = 3 mice), MMAE (*n* = 4 mice), SQL70 (*n* = 4 mice), and SQL70 with SQP22 dosed at 2x (*n* = 5 mice) and 3x (n = 5 mice) molar equivalents of MMAE/dose. **B**, Maximum body weight loss (shown as percentage) was assessed up to Day 7 to determine acute effects on body weight, since RENCA cells induce body weight loss as tumors grow. *P*-values were determined by one-way ANOVA with Bonferroni correction for multiple comparisons, compared to the vehicle group. ANOVA, analysis of variance; MMAE, monomethyl auristatin E; MTD, maximum tolerated dose.

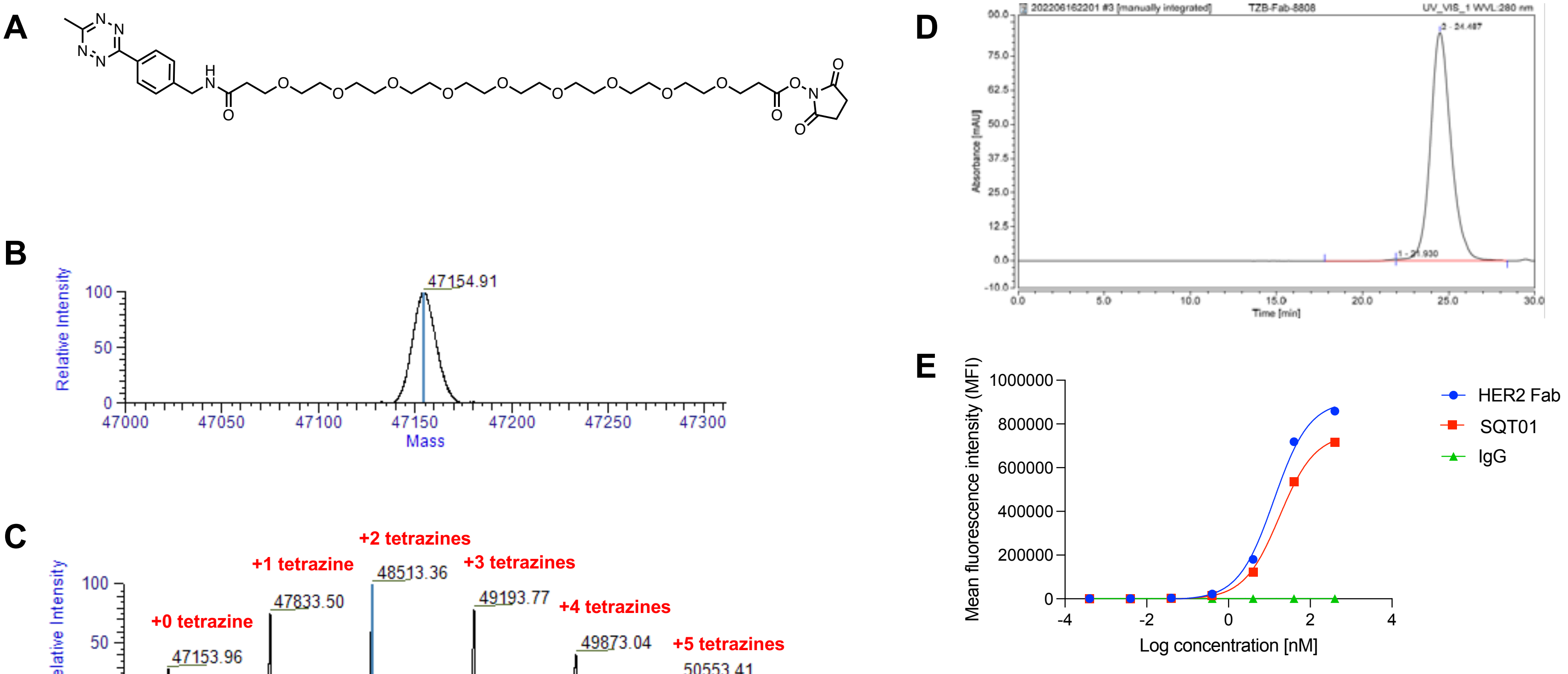


| NCI-N87 cells | | |
|---|---|---|
| Antibody (Ab) ID | max MFI | EC50 (μg/mL) |
| HER2 Fab | 869,554 | 0.5849 |
| SQT01 | 737,310 | 0.8487 |
| IgG | ~ 1,141 | / |

**Figure S4.** Characterization of SQT01. **A**, Structure of tetrazine-PEG9-NHS used to modify the Fab. **B**, Mass spectrum of unmodified Fab. **C**, Mass spectrum of tetrazine-modified Fab (calculated tetrazine-to-antibody ratio was 2.2). **D**, Size exclusion chromatography analysis showing > 99% monomeric species for the Fab-tetrazine conjugate. **E**, Binding of NCI-N87 cells by HER2 Fab, SQT01, and isotype control quantified by flow cytometry. Fab, antigen-binding fragment; HER2, human epidermal growth factor receptor 2.

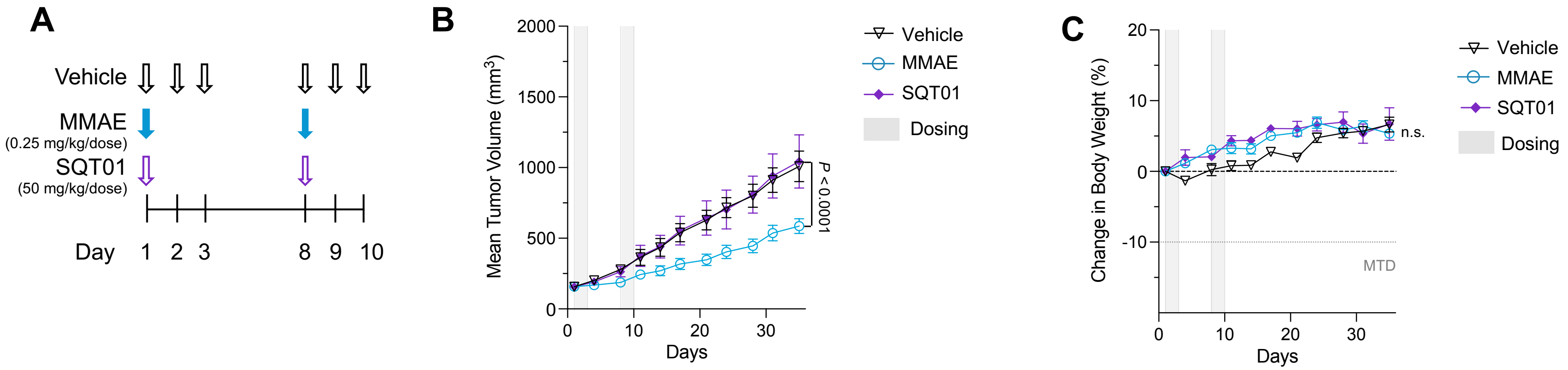


**Figure S5.** SQT01 has no effect on tumor volume or mouse body weight. **A**, Schedule of dosing of agents. **B**, Tumor volumes of NCI-N87 tumors in SCID mice treated with vehicle ($n$ = 10 mice), MMAE ($n$ = 10 mice), and SQT01 ($n$ = 5 mice) in absence of a protodrug. **C**, Percent body weight change for the groups. Shown are means ± SEM. $P$-values were determined by two-way ANOVA with Bonferroni correction for multiple comparisons. MMAE, monomethyl auristatin E; MTD, maximum tolerated dose; n.s., not significant.
